# The genome of a nonphotosynthetic diatom provides insights into the metabolic shift to heterotrophy and constraints on the loss of photosynthesis

**DOI:** 10.1101/2020.05.28.115543

**Authors:** Anastasiia Pendergrass, Wade R. Roberts, Elizabeth C. Ruck, Jeffrey A. Lewis, Andrew J. Alverson

## Abstract

Although most of the tens of thousands of diatom species are obligate photoautotrophs, many mixotrophic species can also use extracellular organic carbon for growth, and a small number of obligate heterotrophs have lost photosynthesis entirely. We sequenced the genome of a nonphotosynthetic diatom, *Nitzschia* sp. strain Nitz4, to determine how carbon metabolism was altered in the wake of this rare and radical trophic shift in diatoms. Like other groups that have lost photosynthesis, the genomic consequences were most evident in the plastid genome, which is exceptionally AT-rich and missing photosynthesis-related genes. The relatively small (27 Mb) nuclear genome did not differ dramatically from photosynthetic diatoms in gene or intron density. Genome-based models suggest that central carbon metabolism, including a central role for the plastid, remains relatively intact in the absence of photosynthesis. All diatom plastids lack an oxidative pentose phosphate pathway (PPP), leaving photosynthesis as the main source of plastid NADPH. Consequently, nonphotosynthetic diatoms lack the primary source of NADPH required for essential biosynthetic pathways that remain in the plastid. Genomic models highlighted similarities between nonphotosynthetic diatoms and apicomplexan parasites for provisioning NADPH in their plastids. The ancestral absence of a plastid PPP might constrain loss of photosynthesis in diatoms compared to Archaeplastida, whose plastid PPP continues to produce reducing cofactors following loss of photosynthesis. Finally, *Nitzschia* possesses a complete β-ketoadipate pathway. Previously known only from fungi and bacteria, this pathway may allow mixotrophic and heterotrophic diatoms to obtain energy through the degradation of abundant plant-derived aromatic compounds.

## Introduction

Microbial eukaryotes (protists) capture the full phylogenetic breadth of eukaryotic diversity and so exhibit a wide range of ecological, life history, and trophic strategies that include parasitism, autotrophy, heterotrophy, and mixotrophy, i.e., the ability to acquire energy through both autotrophic and heterotrophic means [1]. Autotrophic and heterotrophic processes respond differently to temperature, so mixotrophs may shift toward heterotrophic metabolism as the ocean warms in response to climate change [2]. Given the contribution of mixotrophs to the burial of atmospheric carbon on the seafloor (the “biological pump”), this physiological shift could have profound impacts on the structure and function of marine ecosystems [3,4]. Although the bioenergetics and genomic architecture of microbial photosynthesis are fairly well characterized [5], the modes, mechanisms, and range of organic substrates and prey items, and the genomic underpinnings of heterotrophy and mixotrophy are highly varied across lineages and, as a result, more poorly understood [6]. Many protist lineages include a mix of obligately autotrophic, mixotrophic, and obligately heterotrophic species. Studies focused on evolutionary transitions between nutritional strategies in these lineages provide a potentially powerful approach toward understanding the ecological drivers, metabolic properties, and genomic bases of the wide variety of trophic modes in protists.

Diatoms are unicellular algae that are widely distributed throughout marine, freshwater, and terrestrial ecosystems. They are prolific photosynthesizers responsible for some 20% of global net primary production [7,8], and like many other protists [3,9], diatoms are mixotrophic, i.e., they can supplement growth through catabolism of small dissolved organic compounds from the environment. This form of heterotrophy, sometimes described as osmotrophy, might include both passive diffusion (osmosis) and active transport of dissolved organic carbon compounds for catabolism. Diatoms import a variety of carbon substrates for both growth [10,11] and osmoregulation [12–14], though important questions remain about the full range of organic compounds used by diatoms and whether these vary across species and habitats. Heterotrophic growth has been characterized in just a few species [15–17], but the phylogenetic breadth of these taxa suggests that mixotrophy is a widespread and likely ancestral trait in diatoms.

Like many plant and algal lineages that are ancestrally and predominantly photoautotrophic, some diatoms have completely lost their ability to photosynthesize and are obligate heterotrophs, relying exclusively on extracellular carbon for growth [11]. Unlike some species-rich lineages of photoautotrophs, such as flowering plants and red algae that have lost photosynthesis many times [18,19], loss of photosynthesis has occurred rarely in diatoms, with just two known losses that have spawned a small number (≈ 20) of species [20,21]. These so-called “apochloritic” diatoms maintain colorless plastids with highly reduced, AT-rich plastid genomes that are devoid of photosynthesis-related genes [20,22].

Photosynthesis is central to current models of carbon metabolism in diatoms, where carbon metabolic pathways are highly compartmentalized and, in addition, where compartmentalization varies across the small number of species with sequenced genomes [23]. For example, diatom plastids house the Calvin cycle, which is involved in carbon fixation and supplies triose phosphates for a number of anabolic pathways such as biosynthesis of branched-chain amino acids, aromatic amino acids, and fatty acids [23,24]. The lower payoff phase of glycolysis, which generates pyruvate, ATP, and NADH, is located in the mitochondrion [23]. The centric diatom, *Cyclotella nana* (formerly *Thalassiosira pseudonana*), has a full glycolytic pathway in the cytosol, but compartmentalization of glycolysis appears to differ in the raphid pennate diatoms, such as *Phaeodactylum tricornutum*, which lacks a full glycolytic pathway in the cytosol and carries out the lower phase of glycolysis in both the mitochondrion and the plastid [23]. This type of compartmentalization might help diatoms avoid futile cycles, which occur when two metabolic pathways work in opposite directions within the same compartment [e.g., the oxidative pentose phosphate pathway [PPP] vs. Calvin cycle in the plastid; 23]. All this requires careful orchestration of protein targeting within the cell, as most carbon metabolism genes are encoded by the nuclear genome. Differences in the targeting signals of duplicated genes ensure that nuclear-encoded enzymes end up in the correct subcellular compartment [23,25]. Unlike Archaeplastida, diatom plastids lack a PPP, leaving photosystem I as the primary source of endogenous NADPH cofactors for numerous biosynthetic pathways that take place in the diatom plastid [26].

To gain an alternative perspective on carbon metabolism in diatoms, we sequenced the genome of a nonphotosynthetic species from the genus *Nitzschia*. We used the genome to determine the extent to which central carbon metabolism has been restructured in diatoms that no longer photosynthesize. In addition, this rare transition from mixotrophy to obligate heterotrophy could offer new and unbiased insights into the trophic diversity of diatoms more generally. Our analyses revealed a previously unknown carbon metabolic pathway that may allow mixotrophic and heterotrophic diatoms to metabolize plant-derived biomass and, in addition, allowed us to formulate a hypothesis to account for why photosynthesis has been lost infrequently in diatoms.

## Results

To understand the genomic signatures of the transition from mixotrophy to heterotrophy, we collected and isolated nonphotosynthetic *Nitzschia* [20] from a locale known to harbor several species [27]. We generated an annotated genome for one strain, *Nitzschia* sp. strain Nitz4 (Fig 1). *Nitzschia* Nitz4 grew well in near-axenic culture for approximately one year before dying due to size diminution after failing to reproduce sexually. Although we were not able to assign a name to this species, phylogenetic analyses placed it within a clade composed exclusively of nonphotosynthetic *Nitzschia* [20].

**Fig 1.**
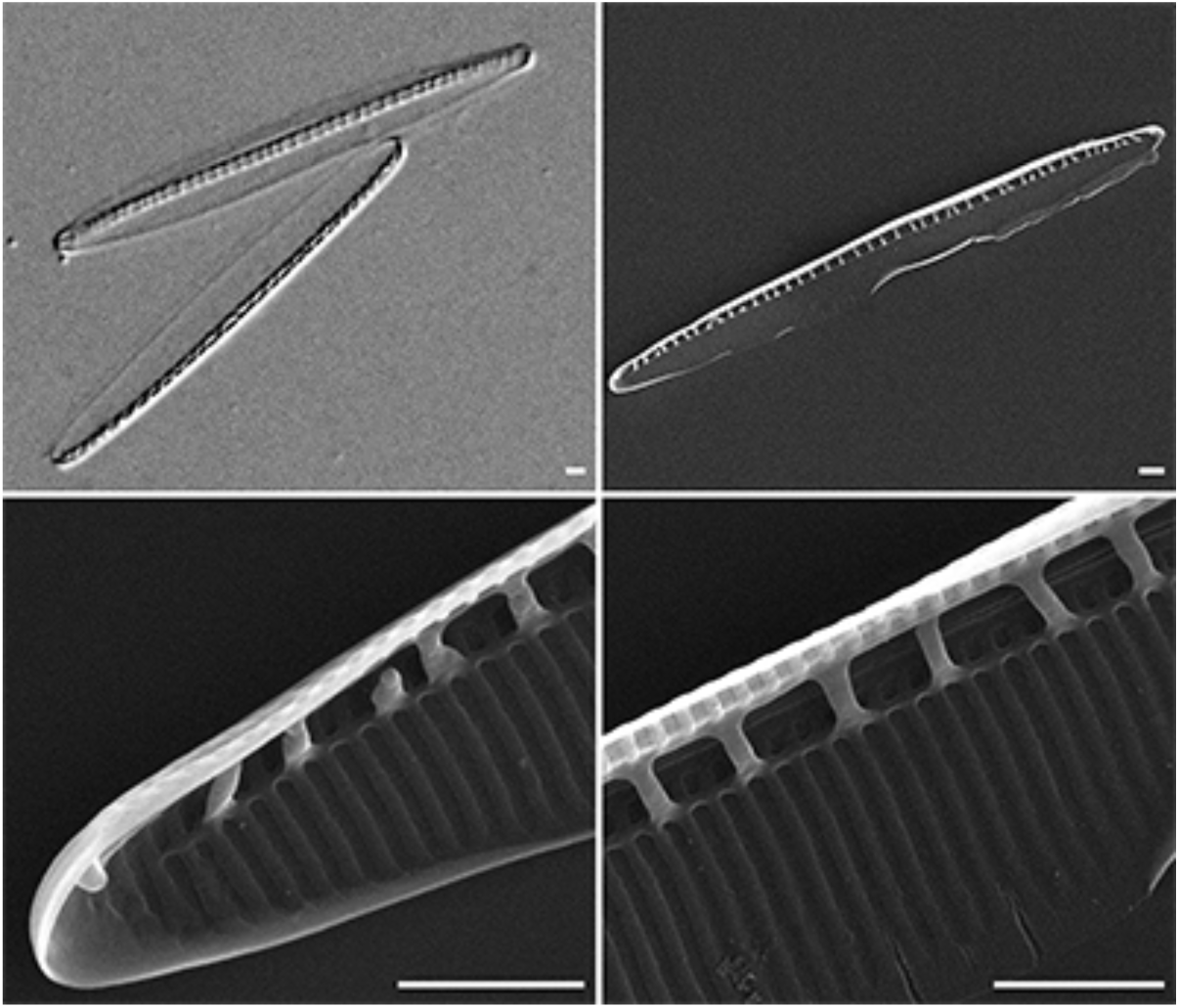
Light and scanning electron micrographs of *Nitzschia* sp. strain Nitz4. Scale bar = 1 micrometer.

### Plastid genome reduction

We examined the fully sequenced plastid genomes of *Nitzschia* Nitz4 [20] and another apochloritic *Nitzschia* [NIES-3581; 22] to understand how loss of photosynthesis impacted the plastid genome. Both plastid genomes revealed a substantial reduction in the amount of intergenic DNA and wholesale loss of genes encoding the photosynthetic apparatus, including those involved in carbon fixation, photosystems I and II, the cytochrome b6f complex, and porphyrin and thiamine metabolism (Table 1). Compared to the plastid genomes of photosynthetic diatoms, which contain a core set of 122 protein and 30 RNA genes [28], the *Nitzschia* Nitz4 plastid genome contains just 95 genes that mostly serve housekeeping functions (e.g., rRNA, tRNA, and ribosomal protein genes). The plastid genome also experienced a substantial downward shift in GC content, from >30% in photosynthetic species to just 22.4% in *Nitzschia* Nitz4 (Table 1), a pattern that is common in streamlined bacterial genomes [29] and may reflect mutational bias [30] or nitrogen frugality [GC base pairs contain more nitrogen than AT base pairs; 31].

**Table 1.** Plastid genome characteristics of five phylogenetically diverse diatoms. Photosynthetic species include *F. cylindrus* and *P. multistriata* and the two model species, *C. nana* and *P. tricornutum*.

|  | *Cyclotella nana* | *Phaeodactylum tricornutum* | *Fragilariopsis cylindrus* | *Nitzschia palea* | *Nitzschia* sp. (Nitz4) |
| --- | --- | --- | --- | --- | --- |
| Genome size | 128,814 | 117,369 | 123,275 | 119,447 | 67,764 |
| GC content (%) | 30.7 | 32.6 | 30.8 | 32.7 | 22.4 |
| Total gene count<sup>1</sup> | 159 | 162 | 141 | 160 | 95 |
| Gene count (photosystem I) | 10 | 10 | 9 | 10 | 0 |
| Gene count (photosystem II) | 18 | 18 | 18 | 18 | 0 |
| Reference | Oudot-Le Secq et al. [32] | Oudot-Le Secq et al. [32] | Zheng et al. [33] | Crowell et al. [34] | Onyshchenko et al. [20] |

<sup>1</sup>Gene count includes rRNAs, tRNAs, and ORFs of unknown function.

### Nuclear genome characteristics

After removal of putative contaminants, a total of 58,141,034 100-bp paired-end sequencing reads were assembled into 992 scaffolds totaling 27.2 Mb in length (Table 2). The genome was sequenced to an average coverage depth of 221 reads. Half of the *Nitzschia* genome was contained in 86 scaffolds, each one longer than 91,515 bp (assembly N50). The average scaffold length was 27,448 bp, and the largest scaffold was 479,884 bp in length. The *Nitzschia* genome is similar in size to other small diatom genomes (Table 2) but is the smallest so far sequenced from Bacillariales, the lineage that includes *Fragilariopsis cylindrus* (61 Mb) and *Pseudo-nitzschia multistriata* (59 Mb) (Table 2). The genomic GC content (48%) is similar to other diatom nuclear genomes (Table 2) and indicates that the exceptional decrease in GC content experienced by the plastid genome (Table 1) did not similarly affect the nuclear (Table 2) or mitochondrial [35] genomes.

**Table 2.**
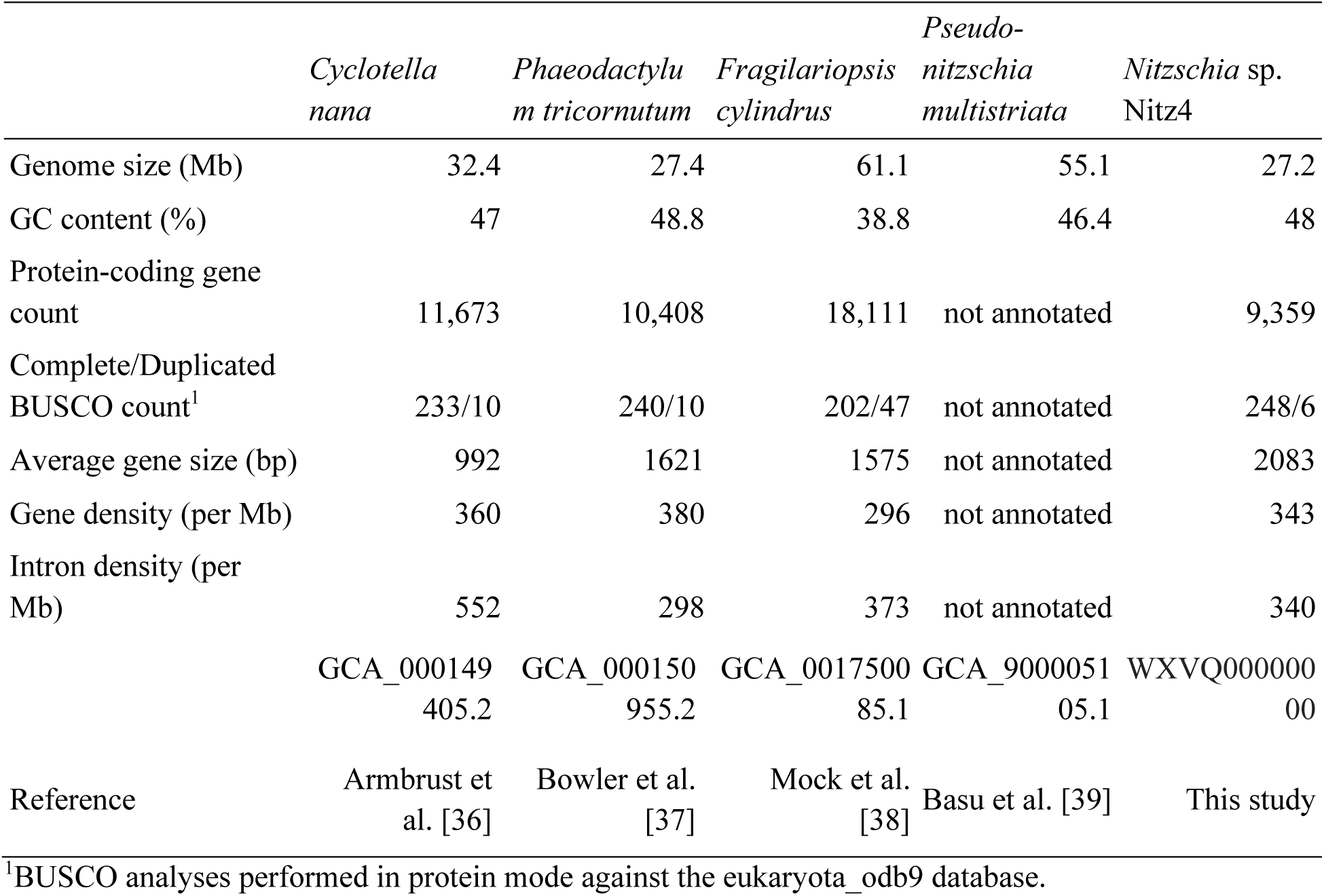
Nuclear genome characteristics of five phylogenetically diverse diatoms. Photosynthetic species include *F. cylindrus* and *P. multistriata* and the two model species, *C. nana* and *P. tricornutum*.

We identified a total of 9,359 protein gene models in the *Nitzschia* Nitz4 nuclear genome. This number increased slightly, to 9,373 coding sequences, when alternative isoforms were included. The vast majority of gene models (9,153/9,359) were supported by transcriptome or protein evidence based on alignments of assembled transcripts (*Nitzschia* Nitz4) and/or proteins (*F. cylindrus*) to the genome. To determine completeness of the genome, we used the Benchmarking Universal Single-Copy Orthologs (BUSCO) pipeline, which characterizes the number of conserved protein-coding genes in a genome [40]. The *Nitzschia* Nitz4 gene models included a total of 254 of 303 eukaryote BUSCOs (Table 2). Although *Nitzschia* Nitz4 has the fewest genes of all diatoms sequenced to date, it has a slightly greater BUSCO count than other diatom genomes (Table 2), indicating that the smaller gene number is not due to incomplete sequencing of the genome. Additional sequencing of Bacillariales genomes is necessary to determine whether the smaller genome size of *Nitzschia* Nitz4 is atypical for the lineage.

Gene models from *C. nana, F. cylindrus, Nitzschia* sp. CCMP2144, and *Nitzschia* Nitz4 clustered into 10,913 orthogroups (S1 Table). A total of 8,068 of the *Nitzschia* Nitz4 genes were assigned to orthogroups, with 13 genes falling into a total of five Nitz4-specific orthogroups (S1 Table). A total of 1,305 genes did not cluster with sequences from other species (S1 Table). Many of these genes (n = 496) had strong matches to other diatom genes but were unplaced by OrthoFinder. The set of 277 unplaced genes with matches to SwissProt were significantly enriched for the heat shock response—perhaps reflecting adaptation of *Nitzschia* Nitz4 to shallow, warm coastal environments—but not for catabolic processes (S2 Table).

We characterized single nucleotide polymorphisms and small structural variants in the genome of *Nitzschia* Nitz4. A total of 77,115 high quality variants were called across the genome, including 62,528 SNPs; 8,393 insertions; and 6,272 deletions (S3 Table). From these variants, the estimated genome-wide heterozygosity was 0.0028 (1 variant per 353 bp), and included 13,446 synonymous and 10,001 nonsynonymous SNPs in protein-coding genes (S3 Table). *Nitzschia* Nitz4 had a higher proportion of nonsynonymous to synonymous substitutions (π_N_/π_S_ = 0.74) than was reported for the diatom, *P. tricornutum* (0.28–0.43) [41,42], and the green alga, *Ostreococcus tauri* (0.2) [43].

### Central carbon metabolism

A key question related to the transition from autotrophy to heterotrophy is how metabolism changes to support fully heterotrophic growth. To address this, we characterized central carbon metabolism in *Nitzschia* Nitz4 to identify genomic changes associated with the switch to heterotrophy. Classically, the major pathways for central carbon utilization in heterotrophs include glycolysis, the oxidative pentose phosphate pathway (PPP), and the Krebs cycle [or tricarboxylic acid (TCA) cycle]. Glycolysis uses glucose (or other sugars that can be converted to glycolytic intermediates) to generate pyruvate and ATP. Pyruvate serves as a key “hub” metabolite that can be used for amino acid biosynthesis, anaerobically fermented to lactate or ethanol, or converted into another hub metabolite, acetyl-CoA. Acetyl-CoA can then be used for fatty acid biosynthesis or can be further oxidized via the Krebs cycle to generate ATP, reducing equivalents (FADH2 and NADH), and certain amino acid precursors. The PPP runs parallel to glycolysis, but unlike glycolysis, is largely anabolic, generating ribose-5-phosphates (a nucleotide precursor) and the key reducing agent NADPH. When the carbon and energy demands of a cell have been met, gluconeogenesis converts pyruvate back into glucose, which can be converted into storage carbohydrates [e.g., chrysolaminarin and trehalose; 44,45] as well as glycolytic intermediates for anabolism [23,46]. Because gluconeogenesis is essentially the reverse of glycolysis and shares many of the same enzymes, flux through each pathway is regulated through the control of specific enzymes driving irreversible reactions (GK, PFK, PK, and FBP).

In diatoms, flux through central metabolism is also controlled through the compartmentalization of different portions of metabolic pathways [23]. For example, in photosynthetic diatoms the oxidative branch of the PPP is only present in the cytoplasm, whereas an incomplete nonoxidative PPP is present in the plastid. As a result, without the NADPH produced by photosystem I, the source of NADPH for biosynthetic pathways that remain localized to the plastid in nonphotosynthetic diatoms is unclear. As a first step in understanding heterotrophic metabolism in *Nitzschia* Nitz4, we examined the genome for differences in the presence or absence, copy number, and enzyme compartmentalization of genes involved in carbon metabolism. We first searched the nuclear genome for genes involved in biosynthesis of the light-harvesting pigments, fucoxanthin and diatoxanthin [47]. We did this by querying the predicted proteins and genomic scaffolds of *Nitzschia* Nitz4 against protein databases for lycopene beta cyclase, violaxanthin de-epoxidase, zeaxanthin epoxidase, and fucoxanthin chlorophyll a/c protein sequences from *C. nana, P. tricornutum*, and *F. cylindrus* [47]. We found no evidence of genes or pseudogenes related to xanthophyll biosynthesis in *Nitzschia* Nitz4, indicating that loss of photosynthesis-related genes was not restricted to the plastid genome.

In general, cytosolic glycolysis in *Nitzschia* Nitz4 does not differ dramatically from other diatoms. For example, despite wholesale loss of plastid genes involved directly in photosynthesis, we detected no major changes in the presence or absence of nuclear genes involved in central carbon metabolism (S4 Table). Like other raphid pennate diatoms, *Nitzschia* Nitz4 is missing cytosolic enolase, which catalyzes the penultimate step of glycolysis, indicating that *Nitzschia* Nitz4 lacks a full cytosolic glycolysis pathway (Fig 2 and S4 Table). *Nitzschia* Nitz4 does have both plastid- and mitochondrial-targeted enolase genes, indicating that the later stages of glycolysis occur in these compartments (Figs 2 and 3; S4 Table). *Nitzschia* Nitz4 has two copies of the gene phosphoglycerate kinase (PGK), a reversible enzyme used in both glycolysis and gluconeogenesis. Enzymes involved in the latter half of glycolysis are localized to the mitochondrion in most diatoms [23,25], but neither copy of the PGK gene in *Nitzschia* Nitz4 contains a mitochondrial targeting signal—instead, one copy is predicted to be plastid-targeted and the other cytosolic (Fig 2 and S4 Table). We verified the absence of a mitochondrial PGK thorough careful manual searching of both the scaffolds and the transcriptome. Lack of a mitochondrial PGK suggests that *Nitzschia* Nitz4 either lacks half of the mitochondrial glycolysis pathway or that one or both of the plastid and cytosolic PGK genes are dual-targeted to the mitochondrion as well. Plastid targeting of PGK may be important for NADPH production in the plastid (see Discussion).

**Fig 2.**
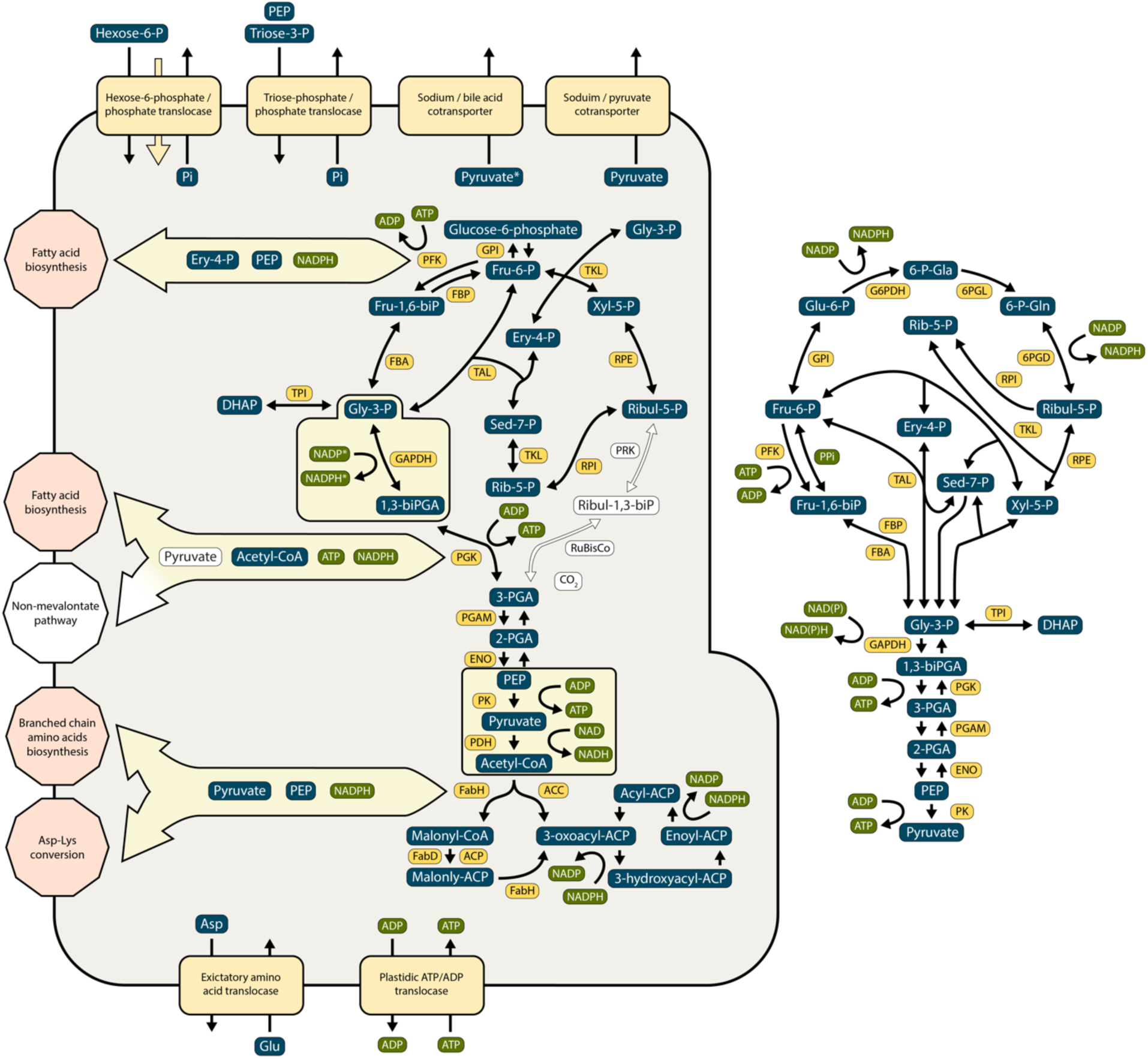
Genome-based model of carbon metabolism in the plastid (green square on left) and cytosol (right) of the nonphotosynthetic diatom, *Nitzschia* sp. Nitz4. Enzymes present in the genome are shown by yellow boxes, and missing enzymes or pathways (arrows) are shown in white. Chemical substrates and products are shown in blue boxes, enzymes encoded by genes present in *Nitzschia* Nitz4 are shown in dark yellow, and reactions that produce or use energy or reducing cofactors are shown in green. Missing genes or pathways are shown in white.

**Fig 3.**
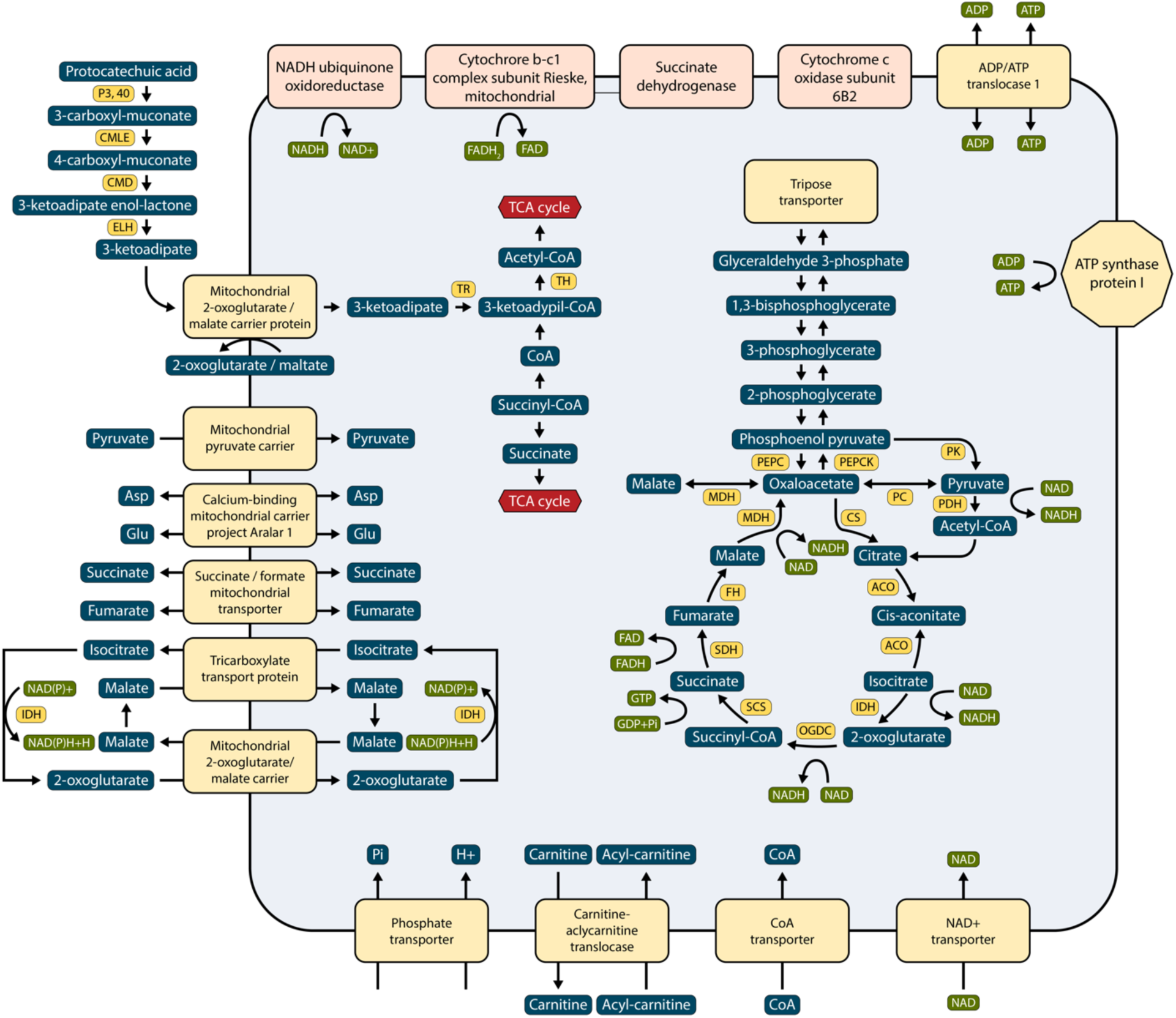
Genome-based model of carbon metabolism in the mitochondria of the nonphotosynthetic diatom, *Nitzschia* sp. Nitz4. Chemical substrates and products are shown in blue boxes, enzymes encoded by genes present in *Nitzschia* Nitz4 are shown in dark yellow, and reactions that produce or use energy or reducing cofactors are shown in green.

Because pyruvate is a key central metabolite that can be used for both catabolism and anabolism, we characterized the presence and localization of “pyruvate hub enzymes” [23] in *Nitzschia* Nitz4 (S4 Table). *Nitzschia* Nitz4 appears to have lost several pyruvate conversion genes, most dramatically in the plastid. For example, the raphid pennate diatoms, *P. tricornutum* and *F. cylindrus*, use pyruvate phosphate dikinase (PPDK) to perform the first step of gluconeogenesis in the plastid, whereas *C. nana* uses a phosphoenolpyruvate synthase (PEPS) that may have been acquired by horizontal gene transfer [23]. *Nitzschia* Nitz4 lacks plastid-localized copies of both enzymes. Additionally, and in contrast to photosynthetic diatoms, *Nitzschia* Nitz4 lacks a plastid-localized pyruvate carboxylase (PC). Together, these results suggest that *Nitzschia* Nitz4 has lost the ability to initiate gluconeogenesis in the plastid and instead relies on mitochondrial-localized enzymes to determine the fate of pyruvate (Figs 2 and 3). The mitochondrial-localized pyruvate hub enzymes of photosynthetic diatoms are mostly present in *Nitzschia* Nitz4 (Fig 3), with the sole exception being malate decarboxylase [or malic enzyme (ME)], which converts malate to pyruvate and CO_2_ [23,25]. In C4 plants, ME is used to increase carbon fixation at low CO_2_ concentrations [48]. If ME performs a similar function in diatoms, then it may be dispensable in nonphotosynthetic species.

Many photosynthesis-related molecules—such as chlorophylls, carotenoids, and plastoquinones—are derivatives of isoprenoids that are produced by the mevalonate pathway (MVA) in the cytosol or, in plastids, by the methylerythritol 4-phosphate pathway (MEP), also known as the non-mevalonate pathway. The absence of an MVA pathway in groups such as apicomplexans, chlorophytes, and red algae suggests that it is not necessary to maintain both MVA and MEP pathways for isoprenoid biosynthesis [49]. The genome of *Nitzschia* sp. Nitz4 has lost its plastid-localized MEP (Fig 2), indicating an inability to synthesize carbon- and NADPH-consuming isoprenoids—the precursors of photosynthetic pigments—in the plastid. A transcriptome study of a closely related nonphotosynthetic *Nitzschia* found a cytosolic MVA but no plastid-localized MEP as well [24].

Finally, we annotated and characterized carbon transporters (S5 Table) and carbohydrate-cleaving enzymes (S6 Table) in *Nitzschia* Nitz4. These analyses suggested that the switch to heterotrophy in *Nitzschia* Nitz4 was not accompanied by expansion or diversification of organic carbon transporters and polysaccharide-digesting enzymes (S1 File).

### β-ketoadipate pathway

One goal of our study was to determine whether apochloritic diatoms, as obligate heterotrophs, are able to exploit organic carbon sources not used by their photosynthetic relatives. To do this, we focused our search on genes and orthogroups specific to *Nitzschia* Nitz4. Although we could not identify or ascribe functions to most of these genes, five of them returned strong but low similarity matches to bacterial-like genes annotated as intradiol ring-cleavage dioxygenase or protocatechuate 3,4-dioxygenase (P3,4O), an enzyme involved in the degradation of aromatic compounds through intradiol ring cleavage of protocatechuic acid (protocatechuate, or PCA), a hydroxylated derivative of benzoate [50]. Additional searches of the nuclear scaffolds revealed three additional P3,4O genes not annotated by Maker (S7 Table). All eight of the P3,4O genes are located on relatively long scaffolds (14–188 kb) without any indication of chimeric- or mis-assembly based on both read depth and consistent read-pair depth. Moreover, the scaffolds also contain genes with strong matches to other diatom genomes (see CoGe genome browser). Taken together, all of the available evidence suggests that these genes are diatom in origin and not the product of bacterial contamination in our conservatively filtered assembly. Later searches revealed homologs for some of the P3,4O genes in other diatom species (S8 Table), providing further support that these are bona fide diatom genes.

In addition to the P3,4O genes, targeted searches of the nuclear scaffolds and annotated gene set revealed that *Nitzschia* Nitz4 has a nearly complete set of enzymes for the β-ketoadipate pathway, including single copies of the 3-carboxymuconate lactonizing enzyme (CMLE), 4-carboxymuconolactone decarboxylase (CMD), and β-ketoadipate enol-lactone hydrolase (ELH) (Fig 3 and S7 Table). All of the genes were transcribed. P3,4O is the first enzyme in the protocatechuate branch of the β-ketoadipate pathway and is responsible for ortho-cleavage of the activated benzene ring between two hydroxyl groups, resulting in β-carboxymuconic acid [50]. Following fission of the benzene ring, the CMLE, CMD, and ELH enzymes catalyze subsequent steps in the pathway, and in the final two steps, β-ketoadipate is converted to the TCA intermediates succinyl-Coenzyme A and acetyl-Coenzyme A [Fig 3; ref. 50]. The enzymes encoding these last two steps—3-ketoadipate:succinyl-CoA transferase (TR) and 3-ketoadipyl-CoA thiolase (TH)—are missing from the *Nitzschia* genome. However, these enzymes are similar in both sequence and predicted function—attachment and removal of CoA—to two mitochondrial-targeted genes that are present in the *Nitzschia* genome, Succinyl-CoA:3-ketoacid CoA transferase (SCOT) and 3-ketoacyl-CoA thiolase (3-KAT), respectively [51,52]. All sequenced diatom genomes, including *Nitzschia* Nitz4, have a putative 4-hydroxybenzoate transporter (PcaK) that reportedly can transport PCA into the cell (S8 Table) [53].

We found evidence of parts of the β-ketoadipate pathway in other sequenced diatom genomes, including a possible complete pathway in the photosynthetic diatom, *Nitzschia* sp. CCMP2144 (S8 Table). If the β-ketoadipate pathway serves an especially important role in *Nitzschia* Nitz4, then we hypothesized that its genome might contain additional gene copies for one or more parts of the pathway. *Nitzschia* Nitz4 has eight copies of the P3,4O gene, compared to four or fewer copies in other diatoms (S8 Table).

## Discussion

### The genomic signatures of loss of photosynthesis in diatoms

Since their origin some 200 Mya, diatoms have become an integral part of marine, freshwater, and terrestrial ecosystems worldwide. Diatoms alone account for 40% of primary productivity in the oceans, making them cornerstones of the “biological pump,” which describes the burial of fixed carbon in the sea floor. Photosynthetic output by diatoms is likely a product of their sheer numerical abundance, species richness, and an apparent competitive advantage in areas of upwelling and high turbulence [54].

Like the genomes of the model photosynthetic diatoms *C. nana* [36] and *P. tricornutum* [37], the *Nitzschia* Nitz4 genome is relatively small and gene dense. Repetitive sequences such as transposable elements are present but in low abundance. Levels of genome-wide heterozygosity are generally <1% in sequenced diatom genomes, ranging from 0.2% in *Pseudo-nitzschia multistriata* [39] to nearly 1% in *C. nana* [36], *P. tricornutum* [37], and *F. cylindrus* [38]. The relatively low observed heterozygosity (<0.2%) and high π_N_/π_S_ (0.74) in *Nitzschia* Nitz4 might reflect a small population size and/or low rate of sexual reproduction [55], hypotheses that cannot be tested with a single genome. Given the small size of the genome, the ability to withstand antibiotic treatments for removal of bacterial DNA contamination, and the abundance of apochloritic *Nitzschia* species in certain locales worldwide [20,56], population-level genome resequencing should provide a feasible path forward for understanding the frequency of sexual reproduction, the effective population sizes, and the extent to which their genomes have been shaped by selection or drift.

Experimental data have shown that mixotrophic diatoms use a variety of carbon sources for growth [10,57]. One goal of our study was to determine whether nonphotosynthetic diatoms have an enhanced capacity for carbon acquisition, through gene family expansions, for example, or by exploiting novel carbon compounds for growth. Aside from losses of genes involved in photosynthesis and pigmentation, we found no major changes in the presence or absence of genes involved in central carbon metabolism and little evidence for expansions of gene families associated with carbon acquisition or carbon metabolism. By examining possible alternative pathways by which *Nitzschia* Nitz4 acquires carbon from the environment, we found evidence for a β-ketoadipate pathway that may allow *Nitzschia* Nitz4 to catabolize plant-derived aromatic compounds. As discussed below, all or parts of this pathway were found in photosynthetic species as well.

Species in the the clade of raphid pennate diatoms, which includes *Nitzschia* Nitz4, are mixotrophic, and mixotrophy is naturally considered as a stepping stone between autotrophy and heterotrophy [58], so genome comparisons to other raphid pennate diatoms might conceal obvious features related to obligate versus facultative heterotrophy. For example, differences between mixotrophs and strict heterotrophs could involve other kinds of changes not investigated here including, for example, adaptive amino acid substitutions, transcriptional regulation, or post-translational modifications of key enzymes. Detailed metabolic flux analysis of a pair of closely related photosynthetic and nonphotosynthetic diatoms performed under a variety of conditions is a logical next step for connecting genomic changes to cellular metabolism.

### A β-ketoadipate pathway in diatoms

We used the *Nitzschia* genome sequence to develop a comprehensive model of central carbon metabolism for nonphotosynthetic diatoms with the goal of understanding how they meet their energy demands in the absence of photosynthesis. We systematically searched for known transporters of organic compounds and other carbon-active enzymes to determine whether *Nitzschia* was enriched for such genes, either through acquisition of novel genes or by duplication of existing ones. By widening our search to include *Nitzschia*-specific genes without known functions, we found one gene—P3,4O—that appeared to be both unique and highly duplicated in *Nitzschia* Nitz4. The major role of P3,4O is degradation of aromatic substrates through the β-ketoadipate pathway. The β-ketoadipate pathway is typically associated with soil-dwelling fungi and eubacteria [50,59,60] and consists of two main branches, one of which converts protocatechuate (PCA) and the other catechol, into β-ketoadipate [50]. Diatoms possess the PCA branch of the pathway. Intermediate steps of the PCA branch differ between fungi and bacteria, and diatoms possess the bacterial-like PCA branch [50].

The β-ketoadipate pathway serves as a funneling pathway for the metabolism of numerous naturally occurring PCA or catechol precursors, all of which feed into the β-ketoadipate pathway through PCA or catechol [50,59]. The precursor molecules are derivatives of aromatic environmental pollutants or natural substrates such as plant-derived lignin [50,59,61,62]. As a key component of plant cell walls, lignin is of particular interest because it is one of the most abundant biopolymers on Earth. Lignin from degraded plant material is a major source of naturally occurring aromatic compounds that feed directly into numerous bacterial and fungal aromatic degradation pathways, including the β-ketoadipate pathway.

The first gene in the β-ketoadipate pathway, P3,4O, catalyzes aromatic ring fission, which is presumably the most difficult step of the pathway because of the high resonance energy and enhanced chemical stability of the carbon ring structure of PCA [59]. It is noteworthy that *Nitzschia* Nitz4 possesses eight copies of the P3,4O gene, whereas other diatoms have four or fewer copies. Expansion of the P3,4O gene family in *Nitzschia* Nitz4 suggests that it might be more efficient than photosynthetic diatoms in cleaving the aromatic ring structure of PCA—the putative bottleneck reaction of this pathway.

The β-ketoadipate pathway is known from fungi and bacteria [50], so its presence in *Nitzschia* Nitz4 was unexpected. Follow-up analyses revealed a putatively complete pathway in the transcriptome of a photosynthetic *Nitzschia* (CCMP2144) and parts of the pathway in other sequenced diatom genomes. These findings suggest that the β-ketoadipate pathway may be an ancestral feature of diatom genomes. That diatoms have a bacterial rather than fungal form of the pathway raises important questions about its origin and spread of this pathway across the tree of life. These questions can best be addressed through comparative phylogenetic analyses of a broader set of diatom, stramenopile, and other pro- and eukaryotic genomes.

Although *Nitzschia* Nitz4 possesses a full and transcribed set of genes for a β-ketoadipate pathway, the genomic prediction—that diatoms with this pathway metabolize an abundant, naturally occurring aromatic compound into products (Succinyl- and Acetyl-CoA) that feed directly into the TCA cycle for energy—requires experimental validation. The similarly surprising discovery of genes for a complete urea cycle in the genome of *Cyclotella nana* [36] was later validated experimentally [63], but in contrast and despite some effort, the genomic evidence in support of C4 photosynthesis in diatoms has not been fully validated [64–66]. It will be important to show, for example, whether diatoms can import PCA and whether the terminal products of the pathway are converted into cellular biomass or, alternatively, whether the pathway is for catabolism of PCA or related compounds. If it is indeed for cellular catabolism, it will be important to determine how much it contributes to the overall carbon budget of both photosynthetic and nonphotosynthetic diatoms. The piecemeal β-ketoadipate pathways in some diatom genomes raise important questions about the functions of these genes, and possibly the pathway overall, across diatoms.

### NADPH limitation and constraints on the loss of photosynthesis in diatoms

Plastids, in all their forms across the eukaryotic tree of life, are the site of essential biosynthetic pathways and, consequently, remain both present and biochemically active following loss of their principal function, photosynthesis [67]. Most essential biosynthetic pathways in the plastid require the reducing agent, NADPH, which is generated through the light-dependent reactions of photosynthesis or the oxidative PPP. The primary-plastid-containing Archaeplastida—including glaucophytes [68], green algae [69,70], land plants [71,72], and red algae [73,74]—have both cytosolic- and plastid-localized PPPs. In plants, the plastid PPP can provide NADPH reductants in nonphotosynthetic cells such as embryos [75–77] and root tissues [78]. In Archaeplastida, the plastid PPP continues to supply NADPH when photosynthesis is lost—something that has occurred at least five times in green algae [e.g., 79,80], nearly a dozen times in angiosperms [18], and possibly on the order of 100 times in red algae [19,81]. Despite their age (ca. 200 My) and estimates of species richness on par with angiosperms and greatly exceeding green and red algae, diatoms are only known to have lost photosynthesis twice [20,21]. Although additional losses may be discovered, the transition from photosynthesis to obligate heterotrophy has occurred rarely in diatoms.

Unlike Archaeplastida, diatoms lack a plastid PPP and therefore rely primarily on photosynthesis for production of NADPH in their plastids [26]. Nevertheless, genomic data from this and other studies [24,82] suggest that the plastids of nonphotosynthetic diatoms continue to be the site of many NADPH-dependent reactions (Fig 2). Without a plastid PPP, however, it is unclear how diatom plastids meet their NADPH demands during periods of prolonged darkness or in species that have permanently lost photosynthesis. Possible enzymatic sources include an NADP-dependent isoform of the glycolytic enzyme GAPDH or conversion of malate to pyruvate by malic enzyme (ME) [23]. In *Nitzschia* Nitz4, NADP-GAPDH is predicted to be targeted to the periplastid compartment, but ME appears to have been lost. It is also possible that ancestrally NADH-producing enzymes have evolved instead to produce NADPH in the plastid, which has likely happened in yeast [83].

In addition to enzymatic production, plastids might acquire NADPH via transport from the cytosol, but transport of large polar molecules like NADPH through the inner plastid membrane may be prohibitive [84]. *Arabidopsis* has a plastid NAD(H) transporter [85], but to our knowledge no NADP(H) transporters have been identified in diatoms. Cells commonly use “shuttle” systems to exchange redox equivalents across intracellular membranes [86]. Although these systems generally shuttle excess reductants from the plastid to the cytosol [84], the triose phosphate/3-phosphoglycerate shuttle is hypothesized to run in reverse to generate NADPH in the relic plastid (apicoplast) of the malaria parasite, *Plasmodium falciparum* [87]. In *P. falciparum*, triose phosphates would be transferred from the cytosol to the apicoplast in exchange for PGA, resulting in the generation of ATP and NADPH in the apicoplast by PGK and NADP-GAPDH (Fig 4). Notably, *Nitzschia* Nitz4 contains a plastid-targeted NADP-GAPDH and two copies of PGK (one cytosolic and one plastid-localized). This differs from photosynthetic diatoms where at least one PGK isoform is generally localized to the mitochondrion [23]. It is possible, therefore, that nonphotosynthetic diatoms also employ a reverse triose phosphate/PGA shuttle to generate plastid NADPH, similar to what has been proposed for apicomplexans (Fig 4). Finally, an apicoplast membrane-targeted NAD(P) transhydrogenase (NTH) was recently discovered in *Plasmodium* [88]. Normally localized to the mitochondrial membrane in eukaryotes, NTH catalyzes (via hydride transfer) the interconversion of NADH and NADPH [Fig 4; ref. 89]. The apicoplast-targeted NTH might provide a novel and necessary source of NADPH to that organelle [90]. Diatoms, including *Nitzschia* Nitz4, have an additional NTH with predicted localization to the plastid membrane (Fig 4), highlighting another way in which nonphotosynthetic diatoms and apicomplexans may have converged upon the same solution to NADPH limitation in their plastids.

**Fig 4.**
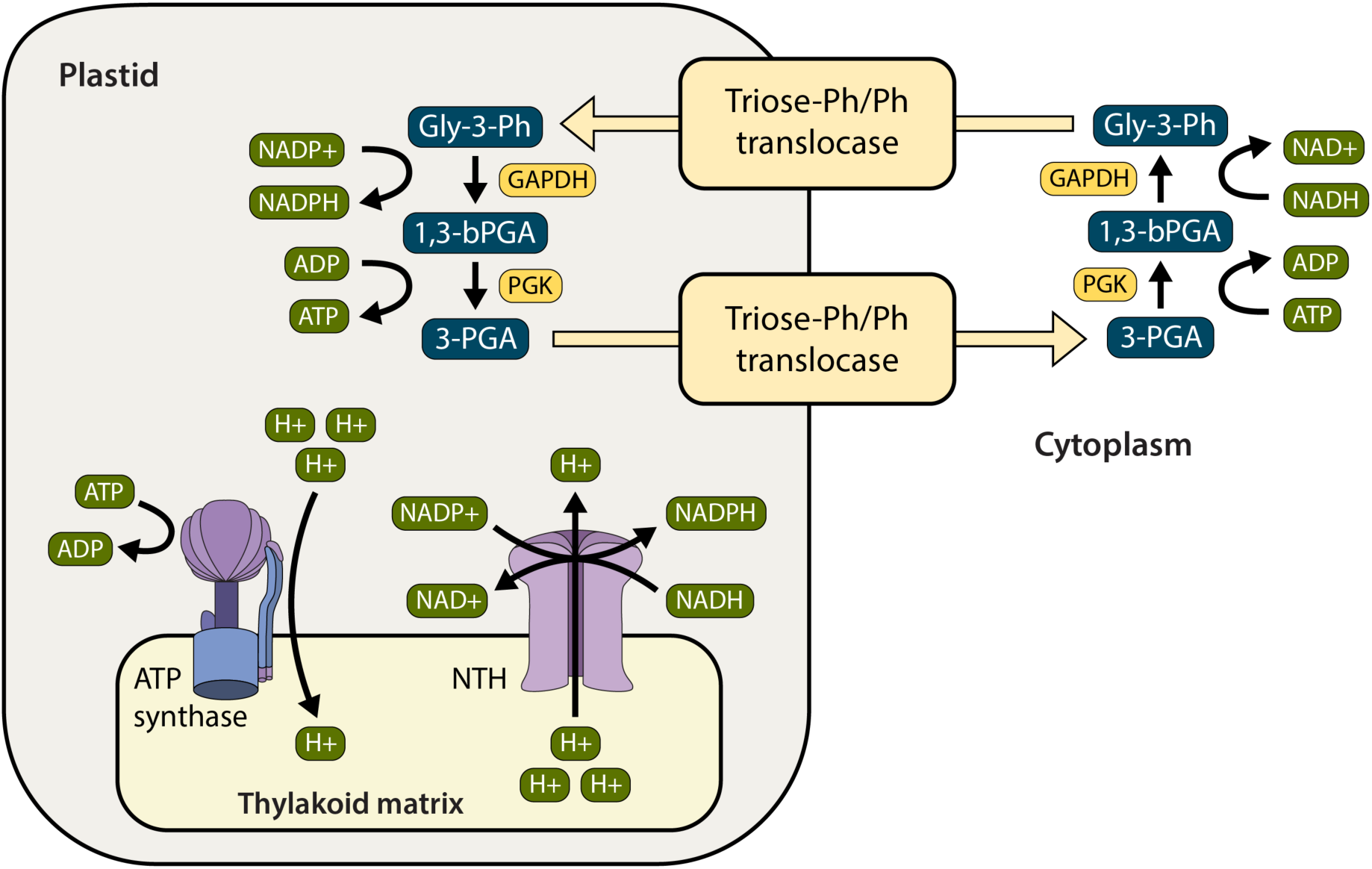
Possible sources of NADPH in the plastid of the nonphotosynthetic diatom, *Nitzschia* sp. Nitz4. The “reverse” triose phosphate/PGA shuttle would exchange cytoplasmic NADH for NADPH, while NAD(P) transhydrogenase (NTH) would use the proton gradient generated by ATP hydrolysis (via ATP synthase) to catalyze interconversion of NADH to NADPH. Chemical substrates and products are shown in blue boxes, and reactions that produce or use protons, energy, or reducing cofactors are shown in green. In addition to ATP synthase and NTH, enzymes encoded by genes present in *Nitzschia* Nitz4 are shown in dark yellow.

Indispensable pathways in diatom plastids include fatty acid biosynthesis, branched chain and aromatic amino acid biosynthesis, and Asp-Lys conversion—all of which have high demands for both carbon and NADPH (Fig 2). In the absence of photosynthesis for carbon fixation and generation of NADPH, and without an ancillary PPP for generation of NADPH, the essential building blocks and reducing cofactors for biosynthetic pathways in the plastid are probably in short supply in *Nitzschia* Nitz4. The need to conserve carbon and NADPH likely drove loss of the isoprenoid-producing MEP from the plastid, which has also been linked to loss of photosynthesis in chrysophytes [91] and euglenophytes [92]. Despite lack of a PPP in the apicoplast [90], *P. falciparum* has not lost its MEP, presumably because it lacks a cytosolic MVA [93], an alternate source of isoprenoid production. Taken together, whether loss of photosynthesis leads to loss of the MEP likely depends on several interrelated factors, including whether the functionally overlapping MVA and MEP pathways [94] were both present in the photosynthetic ancestor and whether, as in Archaeplastida, there is a dedicated plastid PPP. The need to retain expensive pathways in the absence of requisite supply lines may represent an important constraint on loss of photosynthesis in lineages such as diatoms and apicomplexans.

The correlated loss of photosynthesis and the MEP in algae should be tested formally through comparative genomics, whereas the functional hypothesis of limiting carbon and NADPH concentrations in nonphotosynthetic plastids requires comparative biochemistry studies. Whether nonphotosynthetic diatoms have other undescribed mechanisms for provisioning NADPH in their plastids, or whether this is being accomplished by conventional means, remains unclear. Regardless, nonphotosynthetic diatoms offer a promising system for studying these and other outstanding questions related to central carbon metabolism and the evolution of trophic shifts in microbial eukaryotes.

## Conclusions

The goal of our study was to characterize the metabolic shift to heterotrophy from the genome of a nonphotosynthetic diatom, *Nitzschia* sp., one of just a few obligate heterotrophs in a diverse lineage of photoautotrophs that make vast contributions to the global cycling of carbon and oxygen [95]. Aside from loss of the Calvin Cycle and plastid MEP, the genome revealed that central carbon metabolic pathways remained relatively intact following loss of photosynthesis. Moreover, we found no evidence that loss of photosynthesis was accompanied by compensatory gains of novel genes or expansions of genes families associated with carbon acquisition. The genome sequence of *Nitzschia* sp. Nitz4 suggested that nonphotosynthetic diatoms likely rely on the same set of metabolic pathways present in their mixotrophic ancestors. Taken together, these findings suggest that elimination of photosynthesis and the switch from mixotrophy to obligate heterotroph should not be difficult, a hypothesis supported by the experimental conversion of a mixotrophic diatom into a heterotroph through the introduction of a single additional glucose transporter [96]. Nevertheless, given their age (200 My) and species richness (tens of thousands or more species), photosynthesis appears to have been lost less frequently in diatoms than other photosynthetic eukaryotes—a pattern that may reflect diatoms’ dependency on photosynthesis for generation of NADPH in the plastid, which is required to support numerous biosynthetic pathways that persist in the absence of photosynthesis.

The presence of a β-ketoadipate pathway in diatoms expands its known distribution across the tree of life and raises important questions about its origin and function in photosynthetic and nonphotosynthetic diatoms alike. If the genomic predictions are upheld, a β-ketoadipate pathway would allow diatoms to exploit an abundant source of plant-derived organic carbon. The surprising discovery of this pathway in diatoms further underscores the genomic complexity of diatoms and, decades into the genomic era, the ability of genome sequencing to provide an important first glimpse into previously unknown metabolic traits of marine microbes. As for other diatom genomes, we could not ascribe functions to most of the predicted genes in *Nitzschia* Nitz4, potentially masking important aspects of cellular metabolism and highlighting more broadly one of the most difficult and persistent challenges in comparative genomics.

## Materials and Methods

### Collection and Culturing of *Nitzschia* sp. Nitz4

We collected a composite sample on 10 November 2011 from Whiskey Creek, which is located in Dr. Von D. Mizell-Eula Johnson State Park (formerly John U. Lloyd State Park), Dania Beach, Florida, USA (26.081330 latitude, –80.110783 longitude). The sample consisted of near-surface plankton collected with a 10 µM mesh net, submerged sand (1 m and 2 m depth), and nearshore wet (but unsubmerged) sand. We selected for nonphotosynthetic diatoms by storing the sample in the dark at room temperature (21°C) for several days before isolating colorless diatom cells with a Pasteur pipette. Clonal cultures were grown in the dark at 21°C on agar plates made with L1+NPM medium [97,98] and 1% Penicillin–Streptomycin–Neomycin solution (Sigma-Aldrich P4083) to retard bacterial growth.

### DNA and RNA extraction and sequencing

We rinsed cells with L1 medium and removed them from agar plates by pipetting, lightly centrifuged them, and then disrupted them with a MiniBeadbeater-24 (BioSpec Products). We extracted DNA with a Qiagen DNeasy Plant Mini Kit. We sequenced total extracted DNA from a single culture strain, *Nitzschia* sp. Nitz4, with the Illumina HiSeq2000 platform housed at the Beijing Genomics Institute, with a 500-bp library and 90-bp paired-end reads. We extracted total RNA with a Qiagen RNeasy kit and sequenced a 300-bp Illumina TruSeq RNA library using the Illumina HiSeq2000 platform.

### Genome and transcriptome assembly and annotation

A total of 15.4 GB of 100-bp paired-end DNA reads were recovered and used to assemble the nuclear, plastid, and mitochondrial genomes of *Nitzschia*. We used FastQC (ver. 0.11.5) [99] to check reads quality and then used ACE [100] to correct predicted sequencing errors. We subsequently trimmed and filtered the reads with Trimmomatic (ver. 0.32) [101] with settings

“ILLUMINACLIP=<TruSeq_adapters.fa>:2:40:15, LEADING=2 TRAILING=2, SLIDINGWINDOW=4:2, MINLEN=30, TOPHRED64.”

We used SPAdes (ver. 3.12.0) [102] with default parameter settings and k-mer sizes of 21, 33, and 45 to assemble the genome. We then used Blobtools [103] to identify and remove putative contaminant scaffolds. Blobtools uses a combination of GC percentage, taxonomic assignment, and read coverage to identify potential contaminants. The taxonomic assignment of each scaffold was estimated from a DIAMOND BLASTX search [104] with settings “--max-target-seqs 1 --sensitive --evalue 1e-25” against the UniProt Reference Proteomes database (Release 2019_06). We estimated read coverage across the genome by aligning all reads to the scaffolds with BWA-MEM [105]. We used Blobtools to identify and remove scaffolds that met the following criteria: taxonomic assignment to bacteria, archaea, or viruses; low GC percentage indicative of organellar scaffolds; scaffold length < 500 bp and without matches to the UniProt database. After removing contaminant scaffolds, all strictly paired reads mapping to the remaining scaffolds were exported and reassembled with SPAdes as described above. We also used the Kmer Analysis Toolkit [KAT; 106] to identify and remove scaffolds whose length was > 50% aberrant k-mers, which is indicative of a source other than the *Nitzschia* nuclear genome. A total of 401 scaffolds met this criterion and of these, 340 had no hits to the UniProt database and were removed. A total of 55 of the flagged scaffolds had hits to eukaryotic sequences; these were manually searched against the National Center of Biotechnology Information (NCBI) nonredundant (nr) database collection of diatom proteins with BLASTX and removed if they did not fulfill the following criteria: scaffold length < 1000 bp, no protein domain information, and significant hits to hypothetical or predicted proteins only. This resulted in the removal of an additional 31 scaffolds from the assembly. A detailed summary of the genome assembly workflow is available in S2 File.

Due to the fragmented nature of short-read genome assemblies, we used Rascaf [107] and SSPACE [108] to improve contiguity of the final assembly of the relatively small and gene-dense genome of *Nitzschia* Nitz4. Rascaf uses alignments of paired RNA-seq reads to identify new contig connections, using an exon block graph to represent each gene and the underlying contig relationship to determine the likely contig path. RNA-seq read alignments were generated with Minimap2 [109] and inputted to Rascaf for scaffolding (S2 File). The resulting scaffolds were further scaffolded using SSPACE with the filtered set of genomic reads (i.e., ones exported after Blobtools filtering). SSPACE aligns the read pairs and uses the orientation and position of each pair to connect contigs and place them in the correct order (S2 File). Gaps produced from scaffolding were filled with two rounds of GapCloser [110] using both the RNA-seq reads and filtered DNA reads (S2 File). Each stage of the genome assembly was evaluated with QUAST (ver. 5.0.0) [111] and BUSCO (ver. 3.0.2) [40]. BUSCO was run in genome mode separately against both the protist_ensembl and eukaryota_odb9 datasets.

RNA extraction, sequencing, read processing, and assembly of RNA-seq reads followed Parks and Nakov et al. [112]. DNA and RNA sequencing reads are available through NCBI BioProject PRJNA412514. The *Nitzschia* Nitz4 genome is available through NCBI accession WXVQ00000000. The assembled *Nitzschia* Nitz4 and *Nitzschia* Nitz2144 transcriptomes are available from NCBI accessions GIQR00000000 and GIQQ00000000, respectively. A genome browser and gene annotations for *Nitzschia* Nitz4 are available through the Comparative Genomics (CoGe) web platform (https://genomevolution.org/coge/) under genome ID 58203.

### Analysis of heterozygosity

We mapped the trimmed paired-end reads to the genome assembly using BWA-MEM (version 0.7.17-r1188) [105] and used Picard Tools (version 2.17.10) (https://broadinstitute.github.io/picard) to add read group information and mark duplicate read pairs. The resulting BAM file was used as input to the Genome Analysis Toolkit (GATK version 3.5.0) to call variants (SNPs and indels) using the HaplotypeCaller tool [113,114]. SNPs and indels were separately extracted from the resulting VCF file and separately filtered to remove low quality variants. SNPs were filtered as follows: MQ > 40.0, SOR > 3.0, QD < 2.0, FS > 60.0, MQRankSum < −12.5, ReadPosRankSum < −8.0, ReadPosRankSum > 8.0. Indels were filtered as follows: MQ < 40.0, SOR > 10.0, QD < 2.0, FS > 200.0, ReadPosRankSum < - 20.0, ReadPosRankSum > 20.0. The filtered variants were combined and used as “known” variants for base recalibration. We then performed a second round of variant calling with HaplotypeCaller and the recalibrated BAM file. We extracted and filtered SNPs and indels as described above before additionally filtering both variant types by approximate read depth (DP), removing those with depth below the 5^th^ percentile (DP < 125) and above the 95^th^ percentile (DP > 1507). The final filtered SNPs and indels were combined, evaluated using GATK’s VariantEval tool, and annotated using SnpEff (version 4.3t) [115] with the *Nitzschia* Nitz4 gene models.

### Genome annotation

We used the Maker pipeline (ver. 2.31.8) [116] to identify protein-coding genes in the nuclear genome. We used the assembled *Nitzschia* Nitz4 transcriptome (Maker’s expressed sequence tag [EST] evidence) and the predicted proteome of *Fragilariopsis cylindrus* (GCA_001750085.1) (Maker’s protein homology evidence) to inform the gene predictions. RepeatModeler (ver. 2.0) [117] was used for de novo identification and compilation of transposable element families found in the *Nitzschia* Nitz4 genome. The resulting repeat library was searched using BLASTX with settings “-num_descriptions 1 - num_alignments 1 -evalue 1e-10” against the UniProt Reference Proteomes to exclude repeats that had significant protein hits using ProtExcluder (ver. 1.2) [118]. The final custom repeat library was used as input to the Maker annotation pipeline described below.

We used Augustus (ver. 3.2.2) for de novo gene prediction with settings “max_dna_len = 200,000” and “min_contig = 300.” We trained Augustus with the annotated gene set for *Phaeodactylum tricornutum* (ver. 2), which we filtered as follows: (1) remove “hypothetical” and “predicted” proteins, (2) remove all but one splice variant of a gene, (3) remove genes with no introns, and (4) remove genes that overlap with neighboring gene models or the 1000 bp flanking regions of adjacent genes, as these regions are included in the training set along with the enclosed gene. When possible, we annotated untranslated regions (UTRs) by subtracting 5’ and 3’CDS coordinates from the corresponding 5’ and 3’ end coordinates of the associated mRNA sequence. We only retained UTR annotations that extended ≥ 25 bp beyond both the 5’ and 3’ ends of the CDS. The final filtered training set included a total of 726 genes.

We generated a separate set of gene models for UTR training with all of the filtering criteria described above except that we retained intronless genes based on the assumption that intron presence or absence was less relevant for training the UTR annotation parameters. In addition, we retained only those genes with UTRs that were ≥ 40 bp in length. In total, our UTR training set included 531genes. We trained and optimized the Augustus gene prediction parameters (with no UTRs) on the first set of 726 genes and optimized UTR prediction parameters with the second set of 531 genes. We then used these parameters to perform six successive gene predictions within Maker. We assessed each Maker annotation for completeness using BUSCO (ver. 2.0) with the ‘eukaryota_odb9’ and ‘protist_ensembl’ databases [40]. For the five subsequent runs, we followed recommendations of the Maker developers and used a different ab initio gene predictor, SNAP [119], trained with Maker-generated gene models from the previous run. Although the five SNAP-based Maker runs discovered more genes, the number of complete BUSCOs did not increase and the number of genes with Maker annotation edit distance (AED) scores less than 0.5 decreased slightly from 0.98 to 0.97 (S9 Table). We therefore used the gene predictions from the second Maker round (the first SNAP-based round) as the final annotation (S9 Table).

Finally, in our search for genes related to carbon metabolism, we found several genes that were not in the final set of Maker-based annotations. The coding regions of these genes were manually annotated.

### Ortholog clustering and characterization of unplaced genes

We used Orthofinder (ver. 1.1.4) with default parameters to build orthologous clusters from the complete set of predicted proteins from the genomes of *Nitzschia* Nitz4, *Fragilariopsis cylindrus, Phaeodactylum tricornutum, Cyclotella nana*, and the transcriptome of a photosynthetic *Nitzschia* species (strain CCMP2144). A total of 1305 “singleton” genes from *Nitzschia* Nitz4 were not placed into orthogroups by OrthoFinder. We used NCBI BLASTP (-evalue 1e-6) to search these genes against other diatom proteomes to determine whether these were unique to *Nitzschia* Nitz4 or whether they were present in other diatoms but unassigned by OrthoFinder. We broadly characterized the functions of these genes by searching all proteins in the *Nitzschia* Nitz4 genome against the SwissProt database (release 2020_06) with DIAMOND BLASTP (--evalue 1e-6 --very-sensitive --max-target-seqs 1). We then assigned Gene Ontology (GO) terms to each gene using the BLASTP results and the UniProt Retrieve/ID mapping tool (https://www.uniprot.org/uploadlists/). We tested for GO term enrichment within the set of “singleton” genes with the R BioConductor package TopGO using the default “weight01” algorithm and recommended cutoff of *P* < 0.05 [120].

### Characterization of carbon metabolism genes

We performed a thorough manual annotation of primary carbon metabolism genes in the *Nitzschia* Nitz4 genome. We started with a set of core carbon metabolic genes from diatoms [23,24] and used these annotations as search terms to download all related genes, from both diatoms and non-diatoms, from NCBI’s nr database (release 223.0 or 224.0). We then filtered these sets to include only RefSeq accessions. As we proceeded with our characterizations of carbon metabolic pathways, we expanded our searches as necessary to include genes that were predicted to be involved in those pathways. We constructed local BLAST databases of *Nitzschia* Nitz4 genome scaffolds, predicted proteins, and assembled transcripts separately searched the set of sequences for a given carbon metabolism gene against these three *Nitzschia* Nitz4 databases. In most cases, each query had thousands of annotated sequences on GenBank, so if the GenBank sequences for a given gene were, in fact, homologous and accurately annotated, then we expected our searches to return thousands of strong query–subject matches, and this was often the case. We did not consider further any putative carbon genes with weak subject matches (e.g., tens of weak hits vs. the thousands of strong hits in verified, correctly annotated sequences). We extracted the *Nitzschia* Nitz4 subject matches and used BLASTX or BLASTP to search the transcript or predicted protein sequence against NCBI’s nr protein database to verify the annotation. *Nitzschia* Nitz4 subject matches in scaffold regions were checked for overlap with annotated genes, and if the subject match overlapped with an annotated gene, the largest annotated CDS in this region was extracted and searched against the nr protein database with BLASTX to verify the annotation. As necessary, we repeated this procedure for other diatom species that were included in the OrthoFinder analysis.

To identify carbon transporters in *Nitzschia* Nitz4, we searched for genes using the Transporter Classification Database [TCDB; 121] for the following annotations: Channels/Pores, Electrochemical Potential-Driven Transporters, Primary Active Transporters, and Group Translocators. We used the Carbohydrate-Active enZYmes database [CAZy; 122] to identify potential saccharide degradation genes.

### Prediction of protein localization

We used the software programs SignalP [123], ASAFind [124], ChloroP [125], TargetP [126], MitoProt [127], and HECTAR [128] to predict whether proteins were targeted to the plastid, mitochondrion, cytoplasm, or endoplasmic reticulum (ER) for the secretory pathway. Our approach was slightly modified from that of Traller et al. [129]. Proteins that SignalP, ASAFind, and HECTAR predicted to contain a plastid signal peptide were classified as plastid targeted. We placed less weight on ChloroP target predictions, which are optimized for targeting to plastids bound by two membranes rather than the four-membrane plastids found in diatoms. Proteins predicted as mitochondrion-targeted by any two of the TargetP, MitoProt, and HECTAR programs were classified as localized to the mitochondrion. Proteins in the remaining set were classified as ER-targeted if they had ER signal peptides predicted by SignalP and HECTAR. Most of the remaining proteins were cytoplasmic.

## Acknowledgements

This research was carried out by A.P. in partial fulfillment of a Master of Science degree, which was supported by the Fulbright Graduate Student Exchange Program (Ukraine). This research used computational resources available through the Arkansas High Performance Computing Center. We thank Simon Tye for help designing and creating figures, and we thank Ben Lambert for comments on an earlier version of the manuscript.

## Supporting information

**S1 Table**. Summary of orthogroup statistics for the four genomes (*C. nana, P. tricornutum, F. cylindrus*, and *Nitzschia* sp. Nitz4) and one transcriptome (*Nitzschia* CCMP2144) used in the OrthoFinder analysis.

**S2 Table**. Gene ontology enrichment results for the set of genes from *Nitzschia* sp. Nitz4 that were not placed into orthogroups.

**S3 Table**. Summary of variant calling for the nuclear genome of *Nitzschia* sp. Nitz4.

**S4 Table**. Annotation and localization of carbon metabolism genes in *Nitzschia* sp. Nitz4.

**S5 Table**. Annotation of carbon transporters in *Nitzschia* sp. Nitz4.

**S6 Table**. Annotation of carbohydrate-active enzymes in *Nitzschia* sp. Nitz4.

**S7 Table**. Annotation information for genes in the β-ketoadipate pathway of *Nitzschia* sp. Nitz4.

**S8 Table**. β-ketoadipate pathway genes in the sequenced genomes (*C. nana, F. cylindrus, P. tricornutum*, and *P. multiseries*) and transcriptome (*Nitzschia* sp. CCMP2144) of several photosynthetic diatoms.

**S9 Table**. Results of iterative of the genome annotation procedure used to identify and annotate protein-coding genes in the genome of *Nitzschia* sp. Nitz4.

**S1 File**. Summary of carbon transporters and carbohydrate-cleaving enzymes in the *Nitzschia* sp. Nitz4 nuclear genome.

**S2 File**. Assembly and filtering of the *Nitzschia* sp. Nitz4 nuclear genome.

