## Supplementary material for "The genome of a nonphotosynthetic diatom provides insights into the metabolic shift to heterotrophy and constraints on the loss of photosynthesis": S1_File.html

supplementary\_file\_2\_carbohydrate\_and\_transporters.utf8


### Supplementary File 2: Transporters, carbohydrate-cleaving enzymes and expanded gene families

#### Carbon transporters

To find other possible mechanisms of heterotrophic carbon and energy acquisition in *Nitzschia* Nitz4, we characterized genes encoding nutrient transporters and excreted carbohydrate-cleaving enzymes. We also examined whether *Nitzschia* Nitz4 had more copies of genes in these classes compared to photosynthetic diatoms. To do this, we searched genes with annotations of known transporter families in the Transporter Classification Database and compared the number of annotated *Nitzschia* Nitz4 transporter genes for each respective orthogroup to the photosynthetic diatoms *C. nana*, *P. tricornutum*, *F. cylindrus*, and *Nitzschia* Nitz2144. (S5 Table). *Nitzschia* Nitz2144 had more gene copies for multiple orthogroups than the other species, likely due to redundancies (e.g., predicted isoforms) in the transcriptome assembly. Thus, we used *Nitzschia* Nitz2144 to infer presence/absence and not expansion or contraction of gene families in *Nitzschia* Nitz4. The major challenge to this type of analysis is that the substrate specificity for the majority of transporters in diatoms are unknown.

Overall, we found no evidence for major expansions in gene families related to carbon transport in *Nitzschia* Nitz4 compared to other diatoms. In several orthogroups, *Nitzschia* Nitz4 had a few additional paralogs compared to *C. nana*, *P. tricornutum*, and *F. cylindrus*. For example, *Nitzschia* Nitz4 genes had more genes in orthogroup OG0000155 (annotated as ATP-binding cassette transporters with no described function) than other species by a factor of three. *Nitzschia* Nitz4 also had more gene copies (five in total) in orthogroup OG0000212 (similarity with multiple sodium:dicarboxylate symporter/amino acid symporters) compared to *C. nana* (0), *P. tricornutum* (2), and *F. cylindrus* (4). For several other orthogroups, *Nitzschia* Nitz4 had a single additional gene copy compared to other species, e.g., OG0003170 (Amino Acid/Auxin Permease [AAAP] Family), OG0005971 (Lactate Permease Family), and multiple Mitochondrial Carrier Families and ATP-binding Cassette superfamilies.

#### Carbohydrate-active enzymes

We also used the Carbohydrate-Active enZYmes (CAZy) database to determine whether the *Nitzschia* Nitz4 genome has an overrepresentation of genes encoding polysaccharide-digesting enzymes (S6 Table). We focused on two major classes of polysaccharide-digesting enzymes, Glycoside Hydrolases (GHs) and Polysaccharide Lyases (PLs) (S6 Table). For GHs, we grouped roughly 180 specific enzymes based on enzyme functionality. For example, we combined all genes encoding different types of amylase activity (α-amylase, β-amylase, maltotetraose-forming α-amylase) into a single amylase supergroup (S6 Table). There were fewer PL enzymes, so we did not merge these into supergroups (S6 Table). Similar to the organic carbon transport genes, we found no evidence for broad expansion of GHs or PLs in *Nitzschia* Nitz4.

Taken together, these analyses suggest that the switch to heterotrophy in *Nitzschia* Nitz4 was not accompanied by expansion and diversification of organic carbon transporters and polysaccharide-digesting enzymes.
