## Supplementary material for "The genome of a nonphotosynthetic diatom provides insights into the metabolic shift to heterotrophy and constraints on the loss of photosynthesis": S1_Table.html

supplementary\_table\_1\_orthofinder.utf8


### Supplementary Table 1: OrthoFinder analysis

Supplementary Table 2. Summary of orthogroup statistics for the four genomes (*C. nana*, *P. tricornutum*, *F. cylindrus*, and *Nitzschia* sp. Nitz4) and one transcriptome (*Nitzschia* sp. CCMP2144) used in the OrthoFinder analysis.

|  | *Cyclotella nana* | *Phaeodactylum triconutum* | *Fragilariopsis cylindrus* | *Nitzschia* sp. CCMP2144 | *Nitzschia* sp. Nitz4 |
| --- | --- | --- | --- | --- | --- |
| Number of genes | 11673 | 10408 | 18111 | 53477 | 9373 |
| Number of genes in orthogroups | 8746 | 9005 | 11817 | 20372 | 8068 |
| Number of unassigned genes | 2927 | 1403 | 6294 | 33105 | 1305 |
| Percentage of genes in orthogroups | 74.9 | 86.5 | 65.2 | 38.1 | 86.1 |
| Percentage of unassigned genes | 25.1 | 13.5 | 34.8 | 61.9 | 13.9 |
| Number of orthogroups containing species | 7183 | 7653 | 9329 | 10020 | 6594 |
| Percentage of orthogroups containing species | 65.8 | 70.1 | 85.5 | 91.8 | 60.4 |
| Number of species-specific orthogroups | 32 | 8 | 22 | 37 | 5 |
| Number of genes in species-specific orthogroups | 220 | 47 | 136 | 133 | 13 |
| Percentage of genes in species-specific orthogroups | 1.9 | 0.5 | 0.8 | 0.2 | 0.1 |
