## Supplementary material for "The genome of a nonphotosynthetic diatom provides insights into the metabolic shift to heterotrophy and constraints on the loss of photosynthesis": S2_File.html

Supplementary File 2


### Supplementary File 2

#### Genome assembly

##### Software used:

- SPAdes (version 3.12.0)
- Diamond
- BWA
- Samtools
- KAT (K-mer Analysis Toolkit)
- Minimap2
- Rascaf
- SSPACE
- GapCloser (part of SOAPdenovo2)

##### 1. Assemble reads with SPAdes

`spades.py -1 Nitz4_trimmed_R1.fq.gz -2 Nitz4_trimmed_R2.fq.gz -k auto -t 16 -o spades --only-assemble`

**Output files**

- `contigs.fasta`
- `scaffolds.fasta`

##### 2. Filter assembly with Blobtools

###### 2a. Search scaffolds against UniProt Reference Proteomes using BLASTX

`diamond blastx --query scaffolds.fasta --max-target-seqs 1 --sensitive --threads 12 --db uniprot_ref_proteomes.dmnd --evalue 1e-25 --outfmt 6 --out Nitz4_spades.vs.uniprot_ref.mts1.1e25.blastx.out`

###### 2b. Map reads against the scaffolds

`bwa index scaffolds.fasta bwa mem -t 12 scaffolds.fasta Nitz4_trimmed_R1.fq.gz Nitz4_trimmed_R2.fq.gz | samtools sort -@ 12 -o Nitz4_mapped.sorted.bam - samtools index -@ 12 Nitz4_mapped.sorted.bam`

###### 2c. Create Blobtools dataset

`blobtools taxify -f Nitz4_spades.vs.uniprot_ref.mts1.1e25.blastx.out -m uniprot_ref_proteomes.taxids -s 0 -t 2`

`blobtools create -i scaffolds.fasta -b Nitz4_mapped.sorted.bam -t Nitz4_spades.vs.uniprot_ref.mts1.1e25.blastx.taxified.out -o Nitz4`

`blobtools view -i Nitz4.blobDB.json -r superkingdom`

`blobtools plot -i Nitz4.blobDB.json -r superkingdom`

`blobtools plot -i Nitz4.blobDB.json -r phylum`

---

###### Blob plot of inital genome assembly (Superkingdom resolution)

**Figure 1.** Blobology plot of initial genome assembly using all sequencing reads. BLAST hits show contig identity at the superkingdom level.

###### Blob plot of genome assembly (Phylum resolution)

**Figure 2.** Blobology plot of initial genome assembly using all sequencing reads. BLAST hits show contig identity at the phylum level.

###### 2d. Remove the following scaffolds:

1. Scaffolds assigned to bacteria, archaea, or viruses
2. Scaffolds with no hits that are shorter than 500 bp in length
3. Scaffolds with GC content < 35%, indicative of organellar DNA

**Commands:**

`awk '($6 == "Bacteria") || ($6 == "Archaea") || ($6 == "Viruses") || ($6 == "no-hit" && $2 < 500) || ($3 < 0.35) {print $1}' Nitz4.blobDB.table.txt > remove_these.txt`

`blobtools seqfilter -i scaffolds.fasta -l remove_these.txt --invert`

`grep -Fvwf remove_these.txt Nitz4_spades.vs.uniprot_ref.mts1.1e25.blastx.taxified.out > Nitz4_spades.vs.uniprot_ref.mts1.1e25.blastx.taxified.filtered.out`

###### 2e. Plot the filtered dataset

`blobtools create -i scaffolds.filtered.fna -b Nitz4_mapped.sorted.bam -t Nitz4_spades.vs.uniprot_ref.mts1.1e25.blastx.taxified.filtered.out -o Nitz4_filt1`

`blobtools view -i Nitz4_filt1.blobDB.json -r superkingdom`

`blobtools plot -i Nitz4_filt1.blobDB.json -r superkingdom`

`blobtools plot -i Nitz4_filt1.blobDB.json -r phylum`

###### Blob plot of genome assembly after one round of scaffold filtering (Superkingdom resolution)

**Figure 3.** Blobology plot of initial genome assembly filtered according to the criteria in **Remove the following scaffolds** section above. BLAST hits show contig identity at the superkingdom level.

###### Blob plot of genome assembly after one round of scaffold filtering (Phylum resolution)

**Figure 4.** Blobology plot of initial genome assembly filtered according to the criteria in **Remove the following scaffolds** section above. BLAST hits show contig identity at the phylum level.

---

###### 2f. Export the filtered reads

`blobtools bamfilter -b Nitz4_mapped.sorted.bam -e remove_these.txt -o Nitz4_spades_filt1 -n`

**Output files:**

- `Nitz4_spades_filt1.Nitz4_mapped.sorted.bam.1.fa`
- `Nitz4_spades_filt1.Nitz4_mapped.sorted.bam.2.fa`

##### 3. Reassemble the filtered read set with SPAdes

`spades.py -1 Nitz4_spades_filt1.Nitz4_mapped.sorted.bam.1.fa -2 Nitz4_spades_filt1.Nitz4_mapped.sorted.bam.2.fa -k auto -t 16 -o spades_filt1 --only-assemble`

**Output files:**

- `contigs.fasta`
- `scaffolds.fasta`

##### 4. Repeat Blobtools analysis (steps 2a to 2f above)

###### Blob plot of genome assembly after filtering and reassembly (Superkingdom resolution)

**Figure 5.** Blobology plot of genome assembly after one round of scaffold and read filtering and reassembly. BLAST hits show contig identity at the superkingdom level.

###### Blob plot of genome assembly after filtering and reassembly (Phylum resolution)

**Figure 6.** Blobology plot of genome assembly after one round of scaffold and read filtering and reassembly. BLAST hits show contig identity at the phylum level.

##### 5. Use KAT for kmer-based identification of contaminant scaffolds

`kat gcp -o kat-gcp -t 8 Nitz4_spades_filt1.Nitz4_mapped.sorted.bam.1.fa Nitz4_spades_filt1.Nitz4_mapped.sorted.bam.2.fa`

`python3 density.py -x 200 kat-gcp.mx`

**Output files:**

- `kat-gcp.mx`
- `kat-gcp-density.png`

###### 5a. Extract kmers

`kat filter kmer --low_count=25 --high_count=80 Nitz4_spades_filt1.Nitz4_mapped.sorted.bam.1.fa Nitz4_spades_filt1.Nitz4_mapped.sorted.bam.2.fa`

**Output file:** `kat.filter.kmer-in.jf27`

###### 5b. Get scaffolds associated with the filtered k-mers

`kat filter seq --threshold=0.5 --seq=scaffolds.fasta kat.filter.kmer-in.jf27`

**Output file:** `kat.filter.kmer.0.5.in.fasta`

###### Use Blobtools output to manually remove scaffolds with no-hits to the UniProt Reference Proteomes dataset

###### Use BLASTX to manually search the remaining scaffolds against NCBI’s nr database of diatom proteins. Remove scaffolds that meet all the following criteria:

- Scaffold length < 1000 bp
- No protein domains detected
- All hits are to hypothetical or predicted proteins

##### 6. Scaffold the filtered assembly with Rascaf

###### 6a. Map RNA-seq reads against scaffolds with Minimap2

`minimap2 -ax sr scaffolds.filtered.fasta nitz4_trimmed_filtered_RNASEQ_R1.fq.gz nitz4_trimmed_filtered_RNASEQ_R2.fq.gz | samtools sort -@ 12 -o scaffolds.filtered.mapped.bam -`

`samtoojupyls index scaffolds.filtered.mapped.bam`

###### 6b. Use sorted and indexed BAM file as input to Rascaf

`rascaf -b scaffolds.filtered.mapped.bam -f scaffolds.filtered.fasta`

`rascaf-join -r rascaf.out`

**Output file:** `rascaf_scaffold.fa`

##### 7. Scaffold again using SSPACE with the DNA reads

`perl SSPACE_Standard_v3.0.pl -l libraries.txt -s rascaf_scaffold.fa -b nitz4_sspace`

**Output file:** `nitz4_sspace.final.scaffolds.fasta`

##### 8. Extend contigs and fill gaps with GapCloser

###### 8a. Use RNA-seq reads and DNA reads for gap closure

`GapCloser -a nitz4_sspace.final.scaffolds.fasta -b config -o nitz4_sspace.final.scaffolds.filled.fasta -t 16`

###### 8b. Perform a second round of gap closure

`GapCloser -a nitz4_sspace.final.scaffolds.filled.fasta -b config -o nitz4_sspace.final.scaffolds.filled2.fasta -t 16`

##### 9. Make Blobtools plots for the final assembly (steps 2a to 2e above)

###### Blob plot of final genome assembly (Superkingdom resolution)

**Figure 7.** Blobology plot of final genome assembly. BLAST hits show contig identity at the superkingdom level.

###### Blob plot of final genome assembly (Phylum resolution)

**Figure 8.** Blobology plot of final genome assembly. BLAST hits show contig identity at the phylum level.

##### Final assembly file

`nitz4_sspace.final.scaffolds.filled2.fasta`
