## Supplementary material for "The genome of a nonphotosynthetic diatom provides insights into the metabolic shift to heterotrophy and constraints on the loss of photosynthesis": S3_Table.html

supplementary\_table\_3\_heterozygosity.utf8


### Supplementary Table 3: Heterozygosity

Supplementary Table 4. Summary of variant calling for the nuclear genome of *Nitzschia* sp. Nitz4.

|  | *Nitzschia* sp. Nitz4 |
| --- | --- |
| Total variants | 48,874 |
| Number of SNPs | 40,823 |
| Number of insertions | 2,984 |
| Number of deletions | 5,067 |
| Number of transitions (Ts) | 24,758 |
| Number of transversions (Tv) | 16,204 |
| Ts/Tv ratio | 1.54 |
| Heterozygosity estimate (genome-wide) | 0.00169 (or 1 variant per 594 bp) |
| Heterozygosity estimate (exons) | 0.00093 (or 1 variant per 1072 bp) |
| Heterozygosity estimate (introns) | 0.00182 (or 1 variant per 550 bp) |
| Number of synonymous variants | 8,777 |
| Number of non-synonymous variants | 6,559 |
