## Supplementary material for "The genome of a nonphotosynthetic diatom provides insights into the metabolic shift to heterotrophy and constraints on the loss of photosynthesis": S9_Table.html

supplementary\_table\_9\_genome\_annotation.utf8


### Supplementary Table 9: Genome annotation

Supplementary Table 1. Results of iterative of the genome annotation procedure used to identify protein-coding genes in the genome of Nitzschia sp. Nitz4.

| Annotation round | Ab initio algorithm (training round^a^) | Number of gene models | Complete BUSCOs (out of 303^b^) | Complete BUSCOs (out of 215^c^) | Proportion of gene models with AED < 0.5 |
| --- | --- | --- | --- | --- | --- |
| 1 | Maker | 8609 | 253 (83.5%) | 139 (64.7%) | 0.976 |
| **2^d^** | **SNAP (1)** | **9340** | **254 (83.8%)** | **140 (65.1%)** | **0.980** |
| 3 | SNAP (2) | 9387 | 254 (83.8%) | 140 (65.1%) | 0.970 |
| 4 | SNAP (3) | 9394 | 254 (83.8%) | 140 (65.1%) | 0.970 |
| 5 | SNAP (4) | 9396 | 254 (83.8%) | 140 (65.1%) | 0.970 |
| 6 | SNAP (5) | 9395 | 254 (83.8%) | 140 (65.1%) | 0.970 |

**^a^** for annotation rounds 2–6, gene predictions used gene models from the previous round as a training set

**^b^** BUSCO protein mode with `eukaryota_odb9` database

**^c^** BUSCO protein mode with `protist_ensembl` database

**^d^** final annotation
